# Tracking single molecule dynamics in the anesthetized *Drosophila* brain

**DOI:** 10.1101/2020.09.18.303818

**Authors:** Adam D. Hines, Bruno van Swinderen

## Abstract

Super-resolution microscopy provides valuable insight for understanding the nanoscale organization within living tissue, although this method is typically restricted to cultured or dissociated cells. Here, we develop a method to track the mobility of individual proteins in *ex vivo* adult *Drosophila melanogaster* brains, focusing on a key component of the presynaptic release machinery, syntaxin1A. We show that individual syntaxin1A dynamics can be reliably tracked within neurons in the whole fly brain, and that the mobility of syntaxin1A molecules increases following conditional neural stimulation. We then apply this preparation to the problem of general anesthesia, to address how different anesthetics might affect single molecule dynamics in intact brain synapses. We find that propofol, etomidate, and isoflurane significantly impair syntaxin1A mobility, while ketamine and sevoflurane have little effect. Resolving single molecule dynamics in intact fly brains provides a novel approach to link localized molecular effects with systems-level phenomena such as general anesthesia.

**Impact statement:** A new approach to track the mobility of individual molecules using intact fly brains reveals a common presynaptic effect for different intravenous and volatile general anesthetics.

## Introduction

Mechanisms of chemical neurotransmission have become increasingly understood over the past several decades (Baker and Hughson, 2016; Han et al., 2017; Südhof, 2012; Sudhof and Rothman, 2009) and this knowledge has uncovered novel hypotheses for how neurotransmission might be compromised by general anesthetics (Bademosi et al., 2018b; Baumgart et al., 2015; Hemmings et al., 2019, 2005; Humphrey et al., 2007; Karunanithi et al., 2020; Troup et al., 2019; van Swinderen and Kottler, 2014). A key recent advancement aiding our understanding of synaptic function is the development super resolution microscopy, which allows for the visualization of proteins and molecules below the diffraction limit of light (Betzig et al., 2006; Willig et al., 2006). Super-resolution microscopy has provided novel insight on the nanoscale structure and dynamics of key components of the presynaptic release machinery, such as syntaxin1A (Bademosi et al., 2016; Reddy-Alla et al., 2017; Ullrich et al., 2015). Photoactivatable localization microscopy (PALM) (Betzig et al., 2006) with single particle tracking in live cells (Manley et al., 2008), allows molecules to be localized and tracked in a variety of systems to explore macromolecular protein dynamics through time (Manzo and Garcia-Parajo, 2015). This has been made possible by the development of photoconvertible fluorophores such as Eos (McKinney et al., 2009; Zhang et al., 2012), which can be attached to proteins of interest in order to stochastically localize molecules sparsely and thereby study protein nanoscale organization, mobility, and diffusion in cells. To study Eos-tagged proteins, dual color illumination in a total internal reflection (TIRF) (Axelrod, 2001) or highly inclined and laminated optical (HILO) (Tokunaga et al., 2008) sheet configuration is employed to simultaneously record and stochastically photoconvert Eos fluorophores in cultured cells or dissociated neurons (Manzo and Garcia-Parajo, 2015). However, there is comparatively little information on single molecule dynamics in more complex living tissue, such as animal brains.

Recently, we have described single molecule imaging in motor nerve terminals of filleted larvae of the fruit fly *Drosophila melanogaster* (Bademosi et al., 2018a, 2016). In that study we tagged the pre-synaptic protein syntaxin1A (Sx1a) with photoconvertible mEos2 and found that genetic activation of motoneurons resulted in increased mobility of Sx1A in the motor nerve terminals, suggesting increased mobilization of the presynaptic machinery when neurons are activated. Sx1a is necessary for the docking and fusion of neurotransmitter-containing vesicles, and is a component of the SNARE complex along with its binding partners SNAP25 and VAMP2 (Südhof, 2012). Sx1a function is highly conserved in all animals (Bennett et al., 1992; Ferro-Novick and Jahn, 1994; Sudhof and Rizo, 2011), with mutations in the protein often implicated in synaptic communication defects and lethality (Fergestad et al., 2001; Fujiwara et al., 2006; Kofuji et al., 2017; Saifee et al., 1998; Schulze et al., 1995; Vardar et al., 2016). A Sx1a gain-of-function mutation was found to confer resistance to volatile general anesthetics in the nematode *Caenorhabditis elegans* (van Swinderen et al., 1999) as well as *Drosophila* flies (Troup et al., 2019), suggesting a potential presynaptic target mechanism for these drugs. Consistent with this view, electrophysiological recordings from the fly neuromuscular junction reveal decreased quantal release under propofol anesthesia (Karunanithi et al., 2020), and super-resolution microscopy showed that propofol immobilizes Syx1a in nanoclusters that are potentially unavailable to form SNARES (Bademosi et al., 2018b). This mechanism has been hypothesized to explain the well-documented reduction of chemical neurotransmission observed in the presence of some general anesthetics (Bademosi et al., 2018b; Baumgart et al., 2015; Covarrubias et al., 2015; Hemmings et al., 2005; Herring et al., 2011, 2009; Karunanithi et al., 2020; Zalucki et al., 2015), but it remains unclear whether this effect is common to all classes of general anesthetics (i.e., volatile and intravenous), and if it is also evident in central synapses in the brain. In recent electrophysiological work, we have shown that clinically relevant concentrations of the intravenous agent propofol decreases the number active release sites at glutamatergic synapses in the fly larval neuromuscular junction, while a propofol analog has no such effect (Karunanithi et al., 2020). This suggests a failure in the recruitment of synaptic release machinery components under propofol, which would be consistent with our finding that propofol immobilizes syntaxin1A in fly larval synapses (Bademosi et al., 2018b). What remains unknown is if a similar effect is evident in central synapses, which would be the more relevant targets for inducing and/or maintaining general anesthesia.

Here, we adapt super resolution imaging techniques to the extracted adult fly brain and utilize this novel approach to test whether diverse intravenous and volatile general anesthetics might share a common presynaptic mechanism in the central nervous system. We found that, similar to *Drosophila* larval neuromuscular junction (Bademosi et al., 2016), the mobility of Sx1a molecules in the adult brain is increased upon neuronal stimulation, thereby providing a physiologically relevant setting to probe for general anesthetic effects in intact brain tissue. We exposed fly brains to diverse volatile and intravenous general anesthetics, to determine if syntaxin1A mobility is also affected in intact brain tissue by some of these commonly used drugs.

## Results and Discussion

### Imaging syntaxin1A mobility in the adult fly brain

We employed single particle tracking PALM (sptPALM) to image and track individual syntaxin1a (Sx1a) molecules in the *ex vivo* brains of adult *Drosophila* fruit flies (Figure 1 A, Figure 1 – figure supplement 1). Sx1a was tagged on the extracellular c-terminus with the photoconvertible fluorophore mEos2 (McKinney et al., 2009) and expressed pan-neuronally (Bademosi et al., 2016). Brains were mounted onto a glass slide in approximately 10 µL of fresh modified hemolymph-like solution 3.1 (Feng et al., 2004) (HL3.1) and sealed with a square coverslip (Menzel-Gläser, ThermoFisher) rimmed with vacuum grease (Dow Corning) (Figure 1A, lower). Light compression reduced the thickness of the brain from approximately 50 µm to 10 µm, allowing for the imaging of tissue in a HILO configuration while retaining neural circuit architecture (Figure 1C, Figure 1 – figure supplement 2). Spinning disc confocal imaging confirmed mEos2 expression in brain neurons (Figure 1D). When observing the brain at 100x magnification, the point spread function (PSF) overlap of the unconverted green form of mEos2 does not allow for the resolution of individual molecules or structures within the fly brain (Figure 1E). Upon exposure to a low intensity ultraviolet (UV, 405 nm) photoconverting stimulus, stochastically switched red mEos2 molecules can be visualized sparsely (Figure 1F). In order to confirm that we were imaging mEos2 molecules, we compared spot counts in brains that had no UV exposure and saw a significant increase in single molecule detection with photoconversion (Figure 1 – figure supplement 3). At 32 msec exposure time, Sx1a-mEos2 molecules can be seen moving inside of neurons of the fly brain (Video supplement 1). Neural structures in the fly brain become evident after performing a maximum projection of a time series of PALM experiments (Figure 1G), confirming that Sx1a-mEos2 is confined.

**Figure 1.**
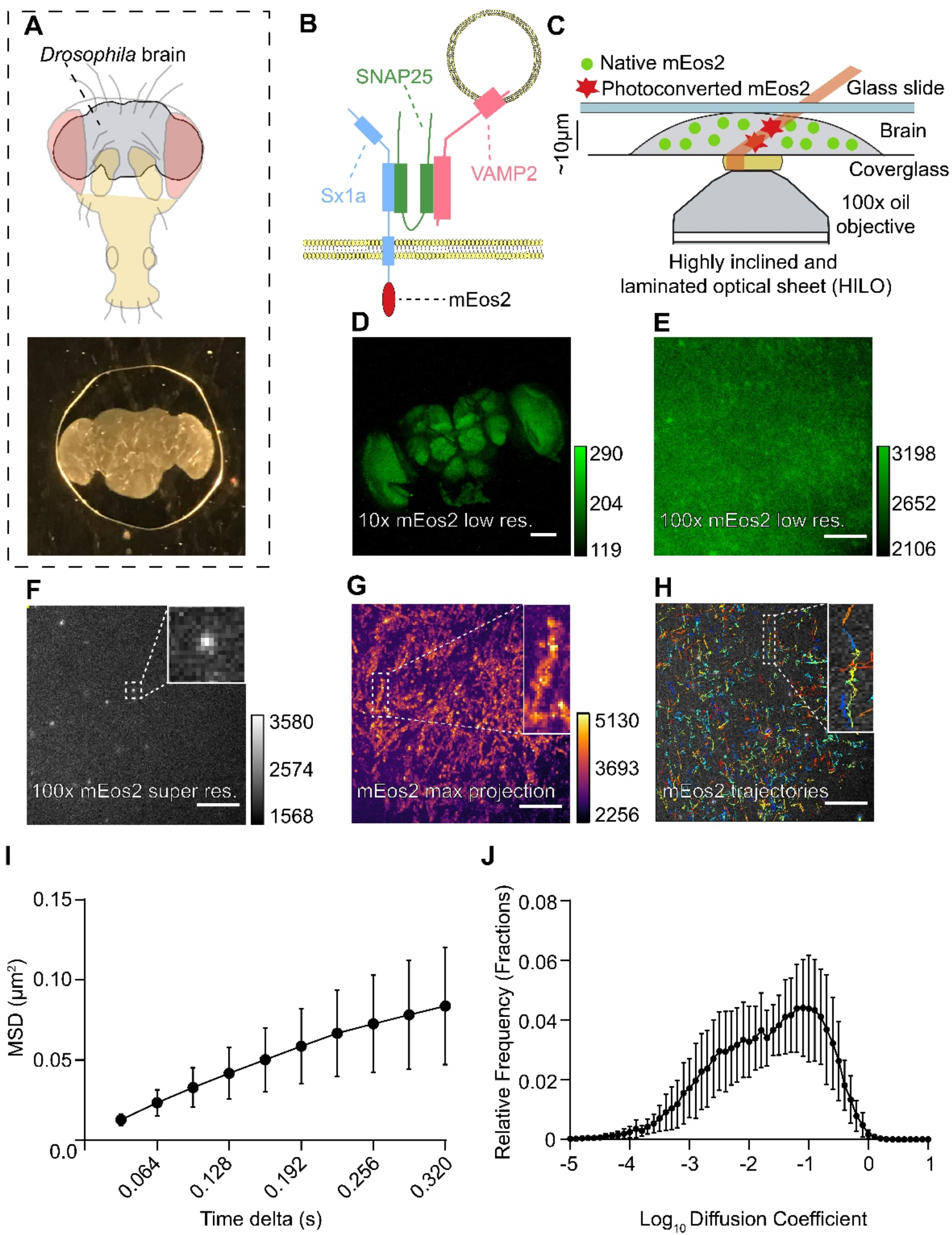
Imaging single syntaxin1a molecules in adult *Drosophila* brains. (A) The brain of adult *Drosophila* fruit flies (top) is dissected and mounted in a HL3.1 buffer and sealed between a glass slide and coverslip (bottom). (B) Schema of the protein of interest being imaged, Syntaxin1a (Sx1a, red), with its SNARE partners SNAP25 (green), and VAMP2 (blue). Sx1a is tagged with the photoconvertible fluorophore mEos2 on the C-terminus. (C) Sx1a-mEos2 expressing brains are imaged under a highly inclined and laminated optical sheet illumination with simultaneous UV-405 nm photoconverting and 561 nm recording lasers. (D) A 10x confocal image showing expression of mEos2 (native non-photoconverted green form) across the entire fly brain (scale bar 100 µm, right calibration scale). (E) Green form of mEos2 expression at 100x magnification, individual molecules cannot be resolved due to point spread function (PSF) overlap (scale bar 5 µm, right calibration scale). (F) Stochastically photoconverting mEos2 with a UV-405 nm laser can resolve single Sx1a-mEos2 molecules using a 561 nm laser without any PSF overlap (scale bar 5 µm, right calibration scale). Inset: digital zoom of one molecule. (G) Neuropil ultrastructure in the fly brain becomes apparent following a maximum intensity projection of all photoconverted mEos2 molecules over 16,000 frames of acquisition (scale bar 5 µm, right calibration scale). Inset: digital zoom of one neuronal compartment. (H) Single particle tracking (SPT) is performed on all detected Sx1a-mEos2 to track individual syntaxin1A molecules. Inset: individual trajectories in different colors. (I & J) Analysis of Sx1a-mEos2 trajectories reveals the mobility of Sx1a-mEos2 by calculating the mean squared displacement (MSD) and diffusion coefficients of single trajectories (n=13 brains, data is ± SD). See Figure 1 - figure supplements 1-4.

In order to characterize the mobility of individual tagged proteins, we performed single particle tracking (SPT, Figure 1 – figure supplement 1) as a post-hoc step to image acquisition. We analyzed on average 2000-3000 individual trajectories of single Sx1a-mEos2 molecules over 16,000 frames (Figure 1H) using the ImageJ software TrackMate (Tinevez et al., 2017) to localize molecules and perform particle tracking (Single Particle Analysis (SPA), see Supplementary file 1). Analysis parameters such as the maximum linking distance for molecules and mean squared displacement (MSD) fitting were set using the SPA software, to perform spot detection and molecule tracking, and then to calculate MSD values (Figure 1I) and molecule diffusion coefficients (Joensuu et al., 2017) (Figure 1J). We confirmed our analysis software by comparing MSD results to the particle tracking software PALM-Tracer, a software used in Metamorph (Molecular Devices) (Figure 1 – figure supplement 4).

To validate the reproducibility of our approach, we compared Sx1a-mEos2 mobility across successive recording sessions from the same brains. We recorded from different brain regions (Figure 2A-C) and from the same brain region (Figure 2F-H). We observed considerable variability in Sx1a-mEos2 mobility across experiments and brain regions (Figure 2D,E), consistent with the large range in MSDs observed in our first dataset (Figure 1I). Crucially, successive recordings from the same region (top right of the central brain, approximately in the lateral protocerebrum) revealed a high level of consistency in the number of localizations, trajectories, and MSD values within a recording site (Figure 2I). This shows that results are repeatable in the same location, but also that some variability in diffusion coefficients exists across experiments in different brains (Figure 2J). Importantly, successive recordings from the same brain region retained a similar number of localizations and trajectories, evident in highly comparable maximum projections of all the single molecule tracks (Figure 2G,H) and the unchanged average spot and trajectory counts (Figure 2 – figure supplement 1). We therefore proceeded with an internally controlled strategy centered on conditional neural activation in sequential recordings from the same location.

**Figure 2.**
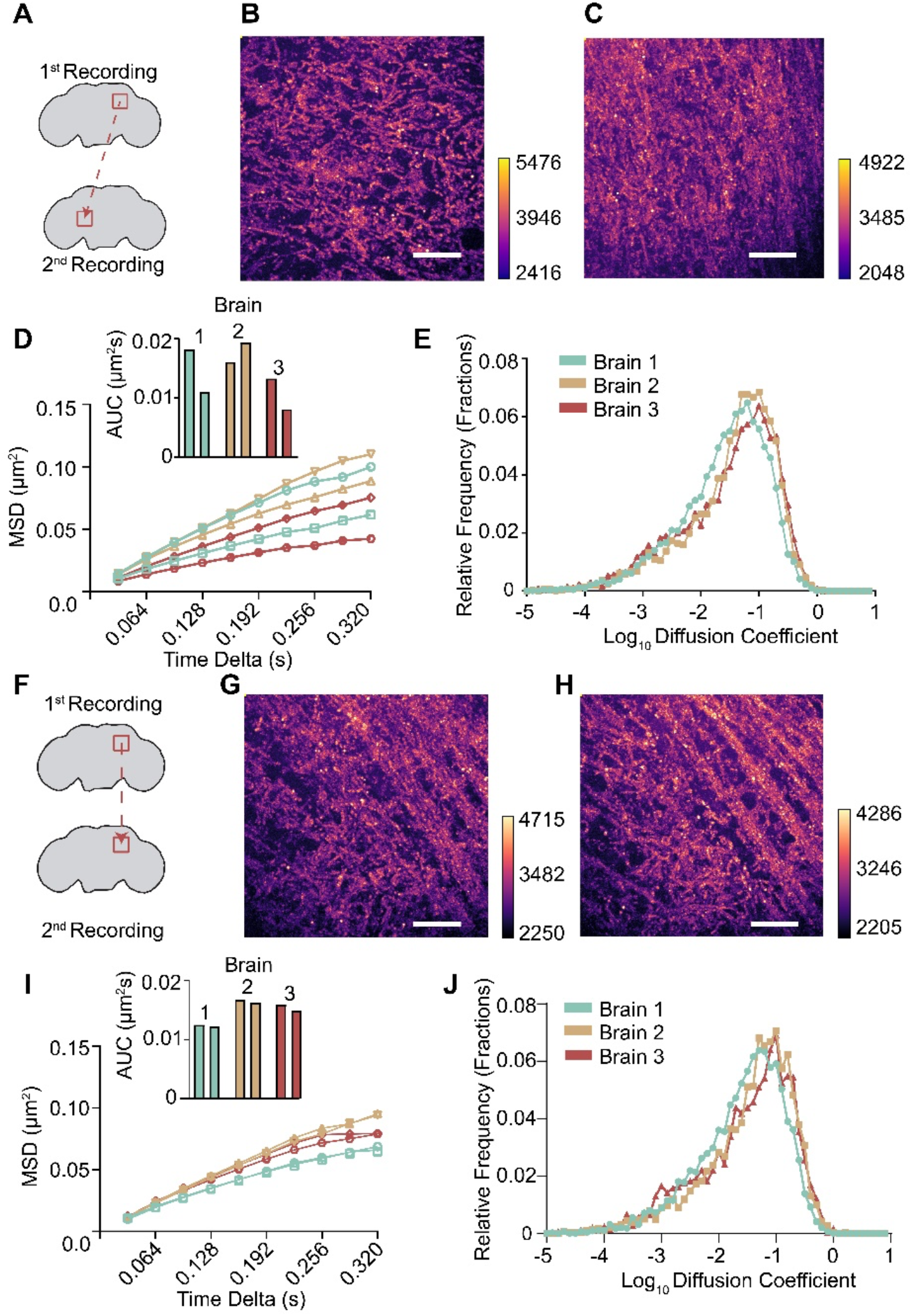
Tracking Sx1a-mEos2 mobility from the same brain region is highly reproducible. (A) Two recordings were taken from different regions in the same brain to establish if differences in Sx1a-mEos2 mobility are observed. (B & C) Maximum stack projections as in Figure 1G for the two distinct brain regions as shown in A reveals different distribution of Sx1a-mEos2 molecules. (D) The MSD and area under the curve (AUC) for 2 successive recordings in 3 separate brains highlights that within a brain there are different levels of Sx1a-mEos2 mobility, resulting in different diffusion coefficient estimates (E) across experiments. (F) Two recordings were taken from the same brain region, to determine if Sx1a-mEos2 mobility was consistent. (G & H) Maximum stack projections for the same brain region recorded twice highlights that the neuronal structure remains the same. (I) When Sx1a-mEos2 in tracked in the same region twice, the MSD and AUC remain consistent, with similar diffusion coefficients (J), providing a framework for internally controlled experiments performed at the same recording site. Scale bars are all 5µm, calibration scales to the right of each. See Figure 2 - figure supplements 1-2.

### Conditional activation of brain neurons increases syntaxin1A mobility

Since the ionic composition of *Drosophila* extracellular fluid buffers varies in different experimental paradigms and can alter neuronal excitability (Feng et al., 2004), we examined the effects of different imaging solutions (Figure 2 – figure supplement 2) and focused on HL3.1 buffer for all subsequent experiments. To ensure that the observed protein mobility was biologically relevant and not an artefact arising from the imaging solution, we performed the same experiment on brains that were first fixed in 4% paraformaldehyde (PFA) and then imaged in HL3.1 solution. Fixing the tissue resulted in a complete loss of Sx1a-mEos2 mobility (Figure 2 – figure supplement 2, Video supplement 2). In addition to this, imaging only HL3.1 solution without any brain tissue revealed highly mobile bright spots that could be localized, but not tracked utilizing our SPA software (Video supplement 3). We next investigated if we could increase Sx1a-mEos2 mobility when we stimulated neurons. In previous work, we have shown that Sx1a-mEos2 mobility increases upon stimulation of larval nerve terminals (Bademosi et al., 2016). To stimulate neurons in the fly brain, we employed a temperature-sensitive *Drosophila* transient receptor potential cation channel 1a (dTrpA1) (Figure 3A), which we expressed under UAS control using the pan-neuronal driver R57c10-Gal4 (Jenett et al., 2012), thereby allowing co-expression with Sx1a-mEos2. Conditional activation of dTrpA1 at 30°C from a baseline of 25°C allowed internally controlled experiments to be performed on the same recording site in the brain (Figure 3B). Thus, all neuronal stimulation data was normalized to the 25°C unstimulated condition at that recording site, thereby controlling for the variability observed across recording sites (Figure 3 – figure supplement 1). When neurons of the fly brain were stimulated, we observed a consistent and significant increase in Sx1a-mEos2 mobility compared to baseline unstimulated conditions (n = 13, p = 0.0002, Wilcoxon test, Figure 3C,D). In contrast, no significant increase in Sx1a-mEos2 mobility was observed at the elevated temperature in control brains that did not express dTrpA1 (Figure 3E,F).

**Figure 3.**
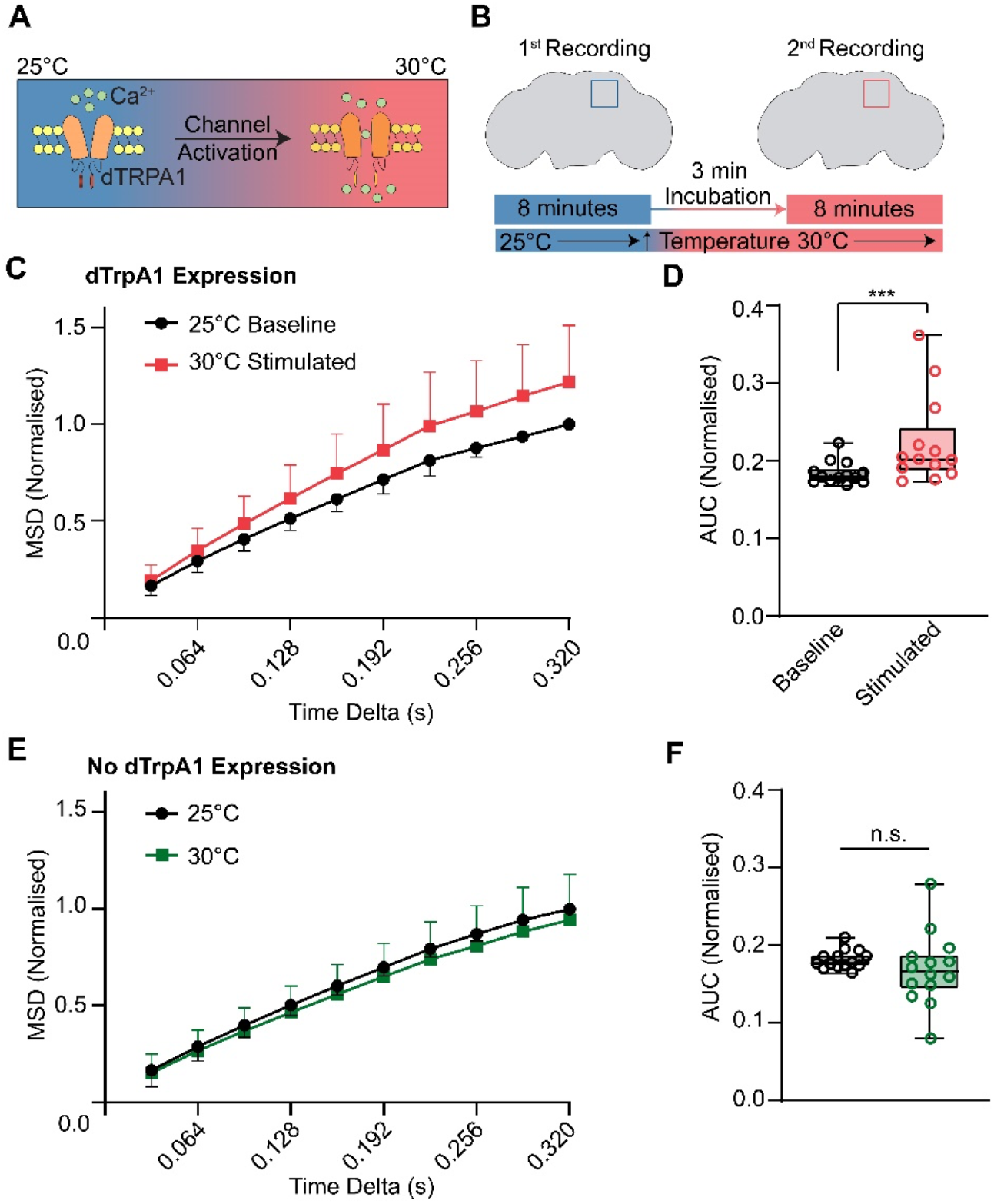
Neuronal stimulation increases the mobility of Sx1a-mEos2. (A) Schematic of the *Drosophila* transient receptor potential cation channel type A1 (dTrpA1) function. At 25°C dTrpA1channels remain closed; increasing ambient temperature to 30°C activates these channels, resulting in Ca2+ influx and neuronal depolarization. (B) To measure the effects of dTrpA1 activation in the fly brain, recordings were taken from the same brain region twice: baseline recording was at 25°C followed by recording at 30°C after increasing incubation temperatures. (C & D) dTrpA1 stimulation of fly neurons increased the mobility of Sx1a-mEos2 molecules compared to baseline. All experiments were normalized to their own internal control at 25°C (n=13, p=0.0002, Wilcoxon test; Area Under Curve (AUC) CI 25°C 0.1747-0.1928, AUC CI 30°C 0.1882-0.2576, data for MSD is ± SD, data for AUC is ± 5-95th percentile). (E & F) In the absence of the R57c10-Gal4 driver, no dTrpA1 was expressed in fly neurons and Sx1a-mEos2 mobility was not increased at 30°C (n=14, p=0.1531, AUC CI 25°C 0.1726 - 0.1885, AUC CI 30°C 0.1425 - 0.1958, Wilcoxon test, data for MSD is ± SD, data for AUC is ± 5-95th percentile). See Figure 3 – figure supplement 1.

### General anesthetics restrict syntaxin1A mobility in brain neurons

Having conditionally increased Sx1a-mEos2 mobility in the fly brain, we next sought to pharmacologically perturb this effect in the same preparation. We have previously shown that the intravenous general anesthetics propofol and etomidate decrease Sx1a-mEos2 mobility in mammalian neurosecretory cells as well as in *Drosophila* motor nerve terminals, by clustering syntaxin1A molecules on the presynaptic membrane (Bademosi et al., 2018b) (Figure 4A). We first investigated if these intravenous general anesthetics also affected Sx1a-mEos2 mobility in the adult *Drosophila* brain, employing our internally controlled strategy. Consistent with our previous findings in other systems (Bademosi et al., 2018b), we found that 3 µM propofol and 8 µM etomidate impaired Sx1a-mEos2 mobility in fly brain neurons (Figure 4B,D). Also consistent with our previous work, a non-anesthetic analog of propofol failed to restrict Sx1a-mEos2 mobility (Bademosi et al., 2018b) (Figure 4 – figure supplement 1). We then proceeded to test other general anesthetics, to see if different categories of drugs also had this immobilizing effect on Sx1a. In contrast to propofol and etomidate, the NMDA-acting sedative ketamine (100 µM) did not affect Sx1a-mEos2 mobility (Figure 4C,D). We next tested two volatile drugs, isoflurane (19 µM), and sevoflurane (38 µM) and found that only the more potent anesthetic isoflurane significantly impaired Sx1A-mEos2 mobility (Figure 4C,D). Combining propofol with sevoflurane, thereby mimicking a typical treatment experienced during surgery in humans (Harris et al., 2006), again significantly impaired Sx1a-mEos2 mobility (Figure 4B,D). Taken together, our results show that Sx1a is highly dynamic in the adult fly brain, with increased mobility following neural stimulation and decreased mobility under general anesthetic exposure. This confirms and expands findings in other model systems (Bademosi et al., 2018b, 2016), and shows that some commonly used intravenous and volatile general anesthetics might share a presynaptic target mechanism centered on Sx1A.

**Figure 4.**
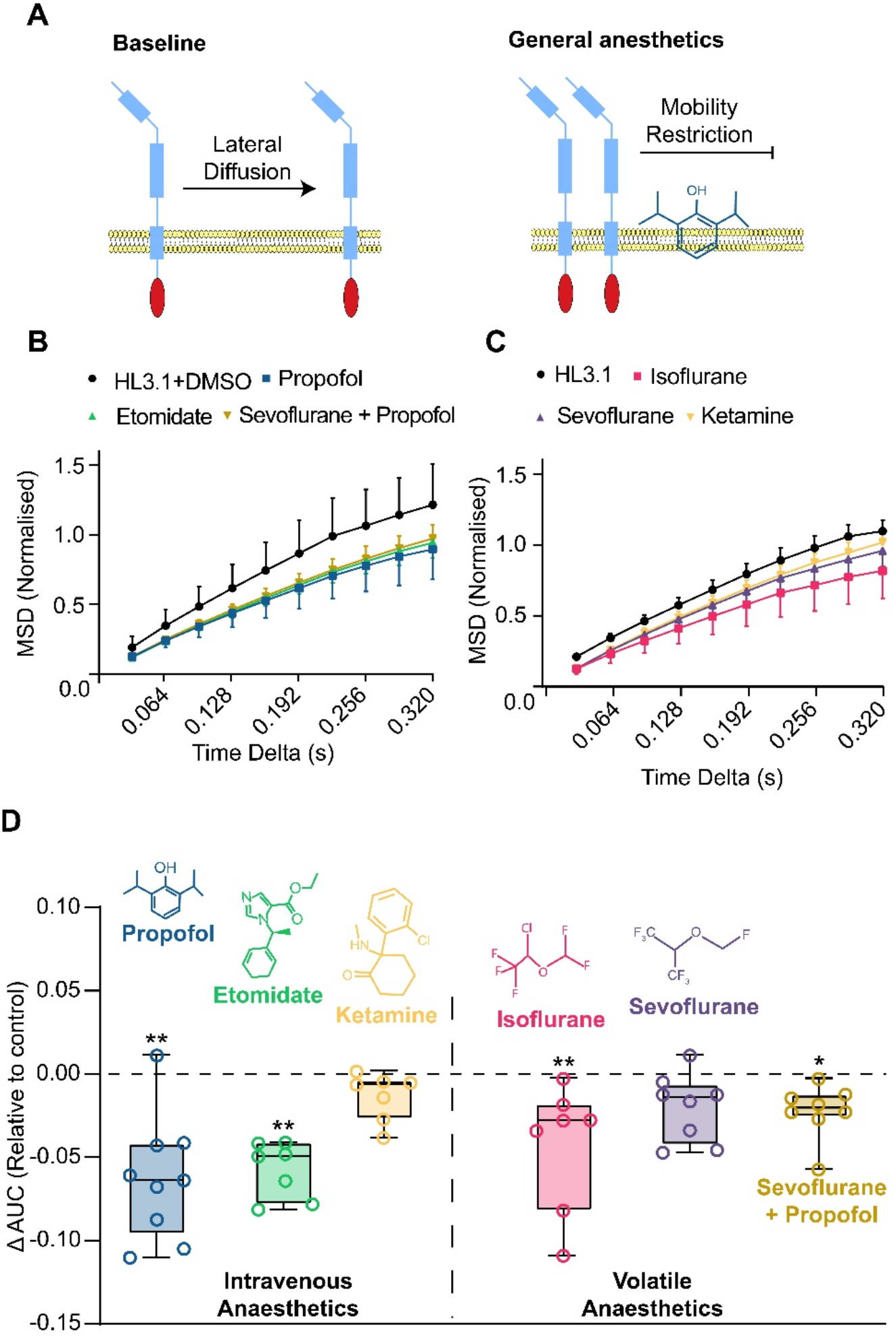
General anesthetics restrict Sx1a-mEos2 mobility in adult *Drosophila* brains. (A) *Top*: Sx1a-mEos2 is able to diffuse laterally across a membrane, but mobility becomes restricted in the presence of propofol *(bottom)*. (B) Normalized MSD curves comparing all anesthetics under stimulation that contained DMSO in the HL3.1. (C) Same as (B) but without DMSO in the solution (MSD is normalized). All data is represented as ± SD. (D) Intravenous and volatile general anesthetics restrict the mobility of Sx1a-mEos2 compared to respective controls (dashed line), ΔAUC values are calculated by subtracting the mean of either HL3.1 or HL3.1+DMSO control from all AUC values for the anesthetic MSD curves (see Figure 3 – figure supplement 1). Both propofol (3 µM) and etomidate (8 µM) significantly reduced Sx1A-mEos2 mobility (ΔAUC) when compared to a HL3.1+DMSO control (propofol n=9, p=0.0009, ΔAUC CI 0.1314 – 0.1886; etomidate n=8, p=0.0055, ΔAUC CI 0.1497 – 0.1811, Kruskal-Wallis test, data is ± 5-95th percentile). Ketamine (100 µM) was unable to restrict Sx1a-mEos2 mobility when compared to a HL3.1 control (n=6, p=0.9924, ΔAUC CI 0.1658 – 0.1923, Kruskal-Wallis test, data is ± 5-95th percentile). The volatile anesthetic isoflurane (19 µM) was able to restrict Sx1a-mEos2 mobility but sevoflurane (38 µM) was not, compared to a HL3.1 control (isoflurane n=9, p=0.0079, ΔAUC CI 0.1140 - 0.1845; sevoflurane n=8, p=0.2672, ΔAUC CI 0.1552-0.1894, Kruskal-Wallis test, data is ± 5-95th percentile). The addition of propofol (3 µM) to sevoflurane significantly restricted Sx1a-mEos2 mobility compared to a HL3.1+DMSO control (n=8, p=0.0108, AUC CI 0.1567 – 0.1837). See Figure 4 – figure supplement 1.

In conclusion, we find that different classes of general anesthetics, such as propofol and isoflurane, have similar effects on the mobility of a presynaptic protein, whereas others such as ketamine and sevoflurane have little effect. General anesthesia has been largely explained as a post-synaptic phenomenon linked to potentiation of inhibitory ion channels (Franks, 2008; Franks and Lieb, 1994; Masiulis et al., 2019). Our results suggest a presynaptic mechanism for some of these drugs, which is consistent with other work done in cell culture and animal models (Bademosi et al., 2018b; Hemmings et al., 2005; Karunanithi et al., 2020; Zalucki et al., 2015). There is increasing evidence that general anesthetics may target mechanisms that influence presynaptic release efficacy, such as mitochondrial function (Morgan et al., 2002), kinesin assembly (Bensel et al., 2017; Woll et al., 2018), channel activation kinetics (Baumgart et al., 2015; Hemmings et al., 2005; Pavel et al., 2020; Zhou et al., 2019) and neurotransmission (Bademosi et al., 2018b; Karunanithi et al., 2020; Troup et al., 2019; Zalucki et al., 2015). More broadly, tracking single particle dynamics in *ex vivo* brains of adult *Drosophila* flies opens a new window into understanding the behavior of individual molecules in intact tissue, to for example help determine which mechanisms are drug-specific and which might reflect a common property of diverse drugs. A major advantage of conducting this work in animal models such as *Drosophila* is the capacity to efficiently test behavioral relevance, as has been shown for resistance-inducing mutations of Sx1a (Troup et al., 2019; van Swinderen & Hines, 2020; Zalucki et al., 2015). Although our results focus on a ubiquitous presynaptic protein expressed in all neurons, the capacity to address circuit-specific questions could be expanded by adapting our method to promoter-driven expression systems such as UAS/Gal4. It will be interesting to apply single molecule imaging to investigate for example if Sx1a is equally compromised at excitatory versus inhibitory synapses, or to examine the individual dynamics of other proteins under general anesthesia, such as receptors in dedicated sleep/wake circuits in the fly brain (Kottler et al., 2013; van Swinderen and Kottler, 2014).

## Author contributions

A.D.H and B.v.S conceptualized the project, developed methodology, and wrote the original draft. A.D.H. performed all experiments. B.v.S. administered the project and acquired funding. Both authors discussed the interpretation of the data and were involved in interpreting results, figure preparation, writing, and editing the manuscript.

## Acknowledgments

We thank Adekunle Bademosi and Merja Joensuu for critical discussions about the work, Rumelo Amor for help with microscopy, and the van Swinderen lab for feedback on the project. This study was funded NHMRC GNT1065715 and GNT1164879 (to BvS) and the Zeiss ELYRA microscope was funded by ARC LIEF LE130100078.

## Materials and Methods

### Fly stocks and rearing conditions

Sx1a-mEos2 transgenic fly lines were generated as previously described (Bademosi et al., 2018a). Briefly, Sx1a cDNA was cloned to include a mEos2 tag by replacing the stop codon of Sx1a with a linker molecule GAGGTACCGCGGGCCCGGGATCCACCG. Whether mEos2 is appropriate for a C- or N-terminal attachment depends on the protein of interest to study. Sx1a-mEos2 flies were injected with phiC31 onto the second chromosome and balanced with curly (Cyo). For dTRPA1 experiments, w^1118^;Sx1a-mEos2/Cyo;+/+ flies were crossed to a w^1118^;+/+;UAS-dTRPA1 line to generate a stable breeding stock with the genotype w^1118^;Sx1a-mEos2/Cyo;UAS-dTRPA1.

*Drosophila melanogaster* fruit flies were reared on standard yeast-sugar-agar food in vials at 22°C with a 12-hour day-night light cycle. w^1118^;Sx1a-mEos2/Cyo;UAS-dTRPA1 transgenic lines were crossed with w^1118^;+/+;R57C10-Gal4 virgin females to generate the w^1118^;Sx1a-mEos2/+;UAS-TrpA1/R57C10-Gal4 flies which were used throughout this study. Flies were raised at 19°C after which point females of the required genotype were collected under brief CO2 exposure and then kept at 19°C on a 12-hour day-night light cycle for 3-5 days before experiments. Keeping the flies at 19°C prevented activation of dTrpA1 channels. The effectiveness of dTrpA1 was confirmed by exposing flies briefly to 30°C, which rapidly induced paralysis (Video supplement 4).

### Imaging Solution

Modified hemolymph-like 3.1 (HL3.1) solution was prepared fresh on the day of an experiment and used both as a dissecting and imaging buffer. HL3.1 consists of 70 mM NaCl, 5 mM KCl, 1.5 mM CaCl2, 2 mM MgCl2, 5 mM HEPES, 115 mM sucrose, 5 mM trehalose, and pH 7.2 with NaHCO3 (Sigma-Aldrich).

### Anesthetics

All anesthetic drugs were diluted into HL3.1 and mixed by vigorous vortexing for ∼1 minute. For intravenous anesthetics, except for ketamine, these were first diluted from stock in dimethyl sulfoxide (DMSO, Sigma-Aldrich D5879-500Ml). Relevant concentrations were determined as previously described (Bademosi et al., 2018b; Zalucki et al., 2015). Volatile anesthetics were taken directly from a stock bottle using a 10 µL Hamilton syringe (Hamilton Company). A fresh preparation of HL3.1 solution with volatile anesthetics was made for each dissection. 3 µL and 6 µL of 100% stock of isoflurane and sevoflurane were diluted into 20 mL of HL3.1 solution respectively (Zalucki et al., 2015). The following anesthetics were used:

- Propofol (2,6-Diisopropylphenol, Sigma-Aldrich D126608-100G)
- Etomidate (Sigma-Aldrich, E6530-10MG)
- Ketamine (Ilium Ketamil, Provet)
- Isoflurane (Henry Schein, 1182097)
- Sevoflurane (Fluorochem, 28523-86-6)

### Dissection of *Drosophila* brains

The brains of 3-5 day old female *Drosophila* flies were removed utilizing a standard dissecting technique (Wu and Luo, 2006) on a Sylgard (Dow Corning) dish after brief anesthesia on a CO2 pad. Using Dumont #5 forceps (Fine Science Tools, 11251-10), heads were removed from the body and placed in HL3.1 solution. The proboscis was then removed to gain access to the inside of the cuticle. Carefully tearing away at the cuticle until the brain is released, the brains were cleared of all tracheal tissue. Dissected brains were then mounted in approximately 10 µL of HL3.1 on a glass slide (Superfrost, ThermoFisher), and sealed shut using a 25mm square coverglass (Menzel-Gläser, ThermoFisher) rimmed with silicone vacuum grease (Dow Corning) with a paintbrush. For fixed brain imaging, brains were dissected as usual and then fixed in 4% paraformaldehyde (PFA) for 40 minutes and then washed in HL3.1 solution. Brains were then mounted in the same manner and imaged.

### Super resolution and photoactivatable localization microscopy

All imaging was performed on a standard Zeiss ELYRA PS.1 microscope fitted with a Zeiss Plan-APOCHROMAT 100x 1.4nA oil immersion objective, a Zeiss FC12 definite focus, and an iXon EMCCD 512×512 pixel camera (Andor, Oxford Instruments). Mounted brains were inverted so that the oil-objective touches the coverslip and the ROI was navigated visually using bright-field illumination. Brains were imaged at a highly inclined and laminated optical sheet (HILO) angle of 47.3° to improve the signal to noise ratio, with a 1.6x lens magnification, in TIRF high power mode. A 570-620 + 750 filter cube was employed to further improve the signal. In order to simultaneously photoconvert native mEos2 and record photoconverted particles, two lasers with 405 nm and 561 nm wavelengths, respectively, were used to perform PALM. Due to a high amount of auto-photoconversion that occurs in the bright-field light from brain dissections, we first allowed the photoconverted particles to bleach for approximately 1 minute without the 405 nm illumination to establish a baseline. Drift was evaluated per brain at this step by finding stable bright spots with the 561 nm illumination, drawing an ROI around the spot, and allowing 3-minutes of continuous recording to occur to see if the spot moved out of the ROI. Brains that drifted were discarded. Drift can often occur due to the movement of recording solution toward the periphery of the coverslip, which can be overcome by sealing the coverslip edges with silicone grease, decreasing the size of the coverslip or increasing the amount of imaging solution. Zeiss Zen 2012 software was used to set the imaging parameters and capture the recordings. For dTRPA1 activation experiments, a Zeiss incubation chamber, Heating Unit XL S, and TempModule S (Zeiss) was utilized to set, change, and monitor recording temperatures. An initial baseline recording at 25°C was taken for all experiments (unless noted otherwise) which was then increased to 30°C to stimulate neurotransmission and perform a second recording at the same location. The power of the UV-405 nm laser was adjusted throughout recordings to maintain the number of stochastically switched mEos2 molecules. A minimum of 16,000 frames were captured at 32 msec exposure time to ensure at least 1,000 Sx1a-mEos2 trajectories were recorded per experiment.

### Data and statistical analysis

All data was analyzed utilizing the free Fiji software TrackMate (Tinevez et al., 2017) written into a custom MATLAB GUI called Single Particle Analysis (SPA, available from www.github.com/AdamDHines). Single Sx1a-mEos2 molecules were localized utilizing a Laplacian of Gaussian (LoG) detection algorithm, median filtering, and subpixel localization with a manually determined threshold value for each recording (Equation 1).

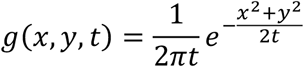

To track single molecules between frames, a linear assignment problem (LAP) algorithm (Jaqaman et al., 2008) was used to link particles by minimizing a cost matrix of distance between detected particles in a frame to every particle in the next frame. A minimum of 6 and a maximum of 1,000 spots per track were included for analysis of the mean squared displacement (MSD), which measures the distance a particle travels from its initial position and is calculated by Equation 2.

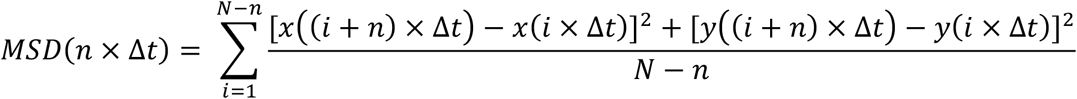

The diffusion coefficient, *D*, was calculated for each MSD curve with linear fits of the first 4 time points using Equation 3.

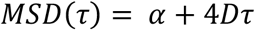

*N* is the number of data points, the offset constant *α* includes the effects of localization error and finite camera exposure, Δ*t* is the time interval between each frame, with *x* and *y* being spatial coordinates for localizations in each image.

For all experiments using thermogenetic stimulation, the peak MSD value for the baseline condition was used to normalize all values of the MSD (Watts et al., 2014) for both unstimulated and stimulated conditions, such that the peak MSD value for the unstimulated condition was set to 1 (Figure 3 – figure supplement 1). The area under the curve (AUC) was measured for each normalized MSD curve. To compare the mean of internally controlled AUC values a Wilcoxon matched signed rank test was used with a significance threshold of p = 0.05. To compare the means of the AUC of different conditions to controls, a Kolmongorov-Smirnov test with a significance threshold of p = 0.05 was used. MSD presented is ± standard deviation (SD) and AUC data is ± 5-95th percentile. 95% Confidence intervals (CI) were calculated around the mean.

## Data and Code Availability

The datasets and code supporting the current study will be made available on a public database upon publication.

## Supplemental information

**Figure 1 – figure supplement 1:**
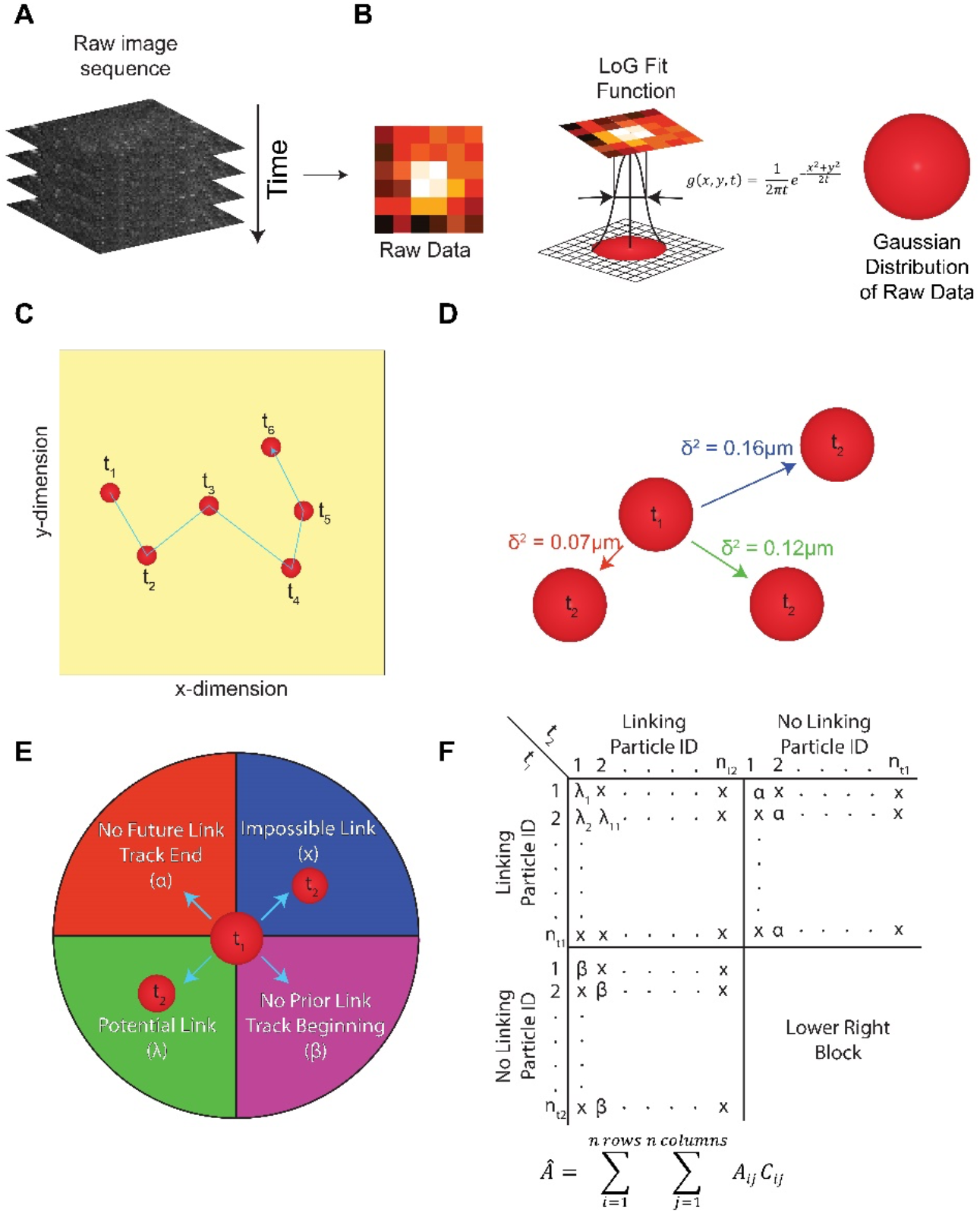
Single particle tracking and photoactivatable localisation microscopy. (A-B) Raw image sequences from photoactivatable localisation microscopy (A) are processed with a Laplacian of Gaussian convolution filter (B) for automatic spot detection to derive centroids of single Sx1a-mEos2 particles. (C-D) Schematic of tracking particles in a 2D-sample over time, with links between frames determined based on the relative distance (δ, distance) of a single particle to every other particles (D) from one frame to the next. (E) Particle tracking is solved using a linear assignment problem cost matrix, where the cost is the relative distance of a particle in frame *n* to every other particle in frame *n+1*. A particle in frame *n* can have one of four outcomes based on the localisation in the proceeding frame. A particle has a potential link (λ) to another particle based on a maximum linking distance which if a particle in the proceeding frame exceeds becomes an impossible link (x). To avoid linking potentially unrelated molecules, it is important to keep stochastic switching of fluorophores light such that molecules detection is sparse. The threshold for the maximum linking distance depends on a variety of factors, including the exposure time of the imaging and the relative speed of the molecule, and if it is membrane bound or cytoplasmic. A particle can also either be the start or the end of a trajectory, and a higher cost value is employed to determine if a particle should be linked to another particle or not (α & β). (F) Example of the cost matrix utilised to solve single particle tracking. The matrix is solved for least cost to link particles and determine if a trajectory is at its beginning or its end (Adapted from Jaqaman *et al*. 2008 (Jaqaman et al., 2008)).

**Figure 1 – figure supplement 2:**
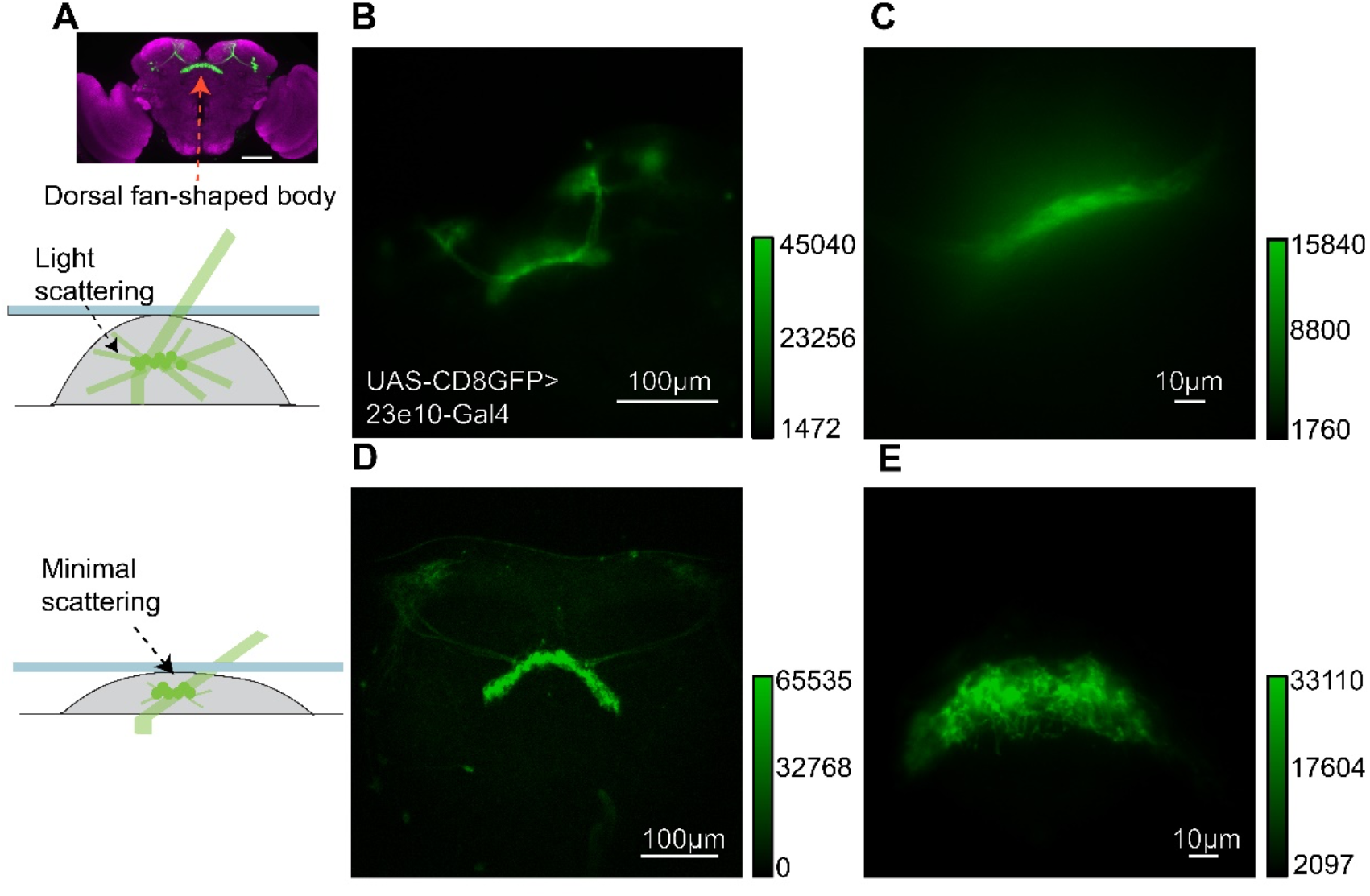
Internal brain structures remain intact in a compressed preparation. (A) Bruchpilot (nc82) stain of the *Drosophila* brain with a UAS-CD8GFP expressed in the dorsal fan-shaped body using the R23e10-Gal4 driver (Jenett et al., 2012) (*Top*, scale bar 100 µm). When the fly brain is not compressed, light scattering under a highly inclined and laminated optic (HILO) sheet setting decreases the resolution of imaged structures (*middle*). When the brain is lightly compressed, the scattering interferes less, and the structure is more resolved (*bottom*). (B-C) 10x and 63x oil magnification, respectively, of UAS-CD8GFP>R23e10-Gal4 imaging in a standard uncompressed preparation. (D-E) 10x and 63x oil magnification, respectively, of UAS-CD8GFP>R23e10-Gal4 imaging in a compressed preparation, revealing that neural architecture in the fly brain remains intact. Bars to the right of graph in (B-E) are calibration scales.

**Figure 1 – figure supplement 3:**
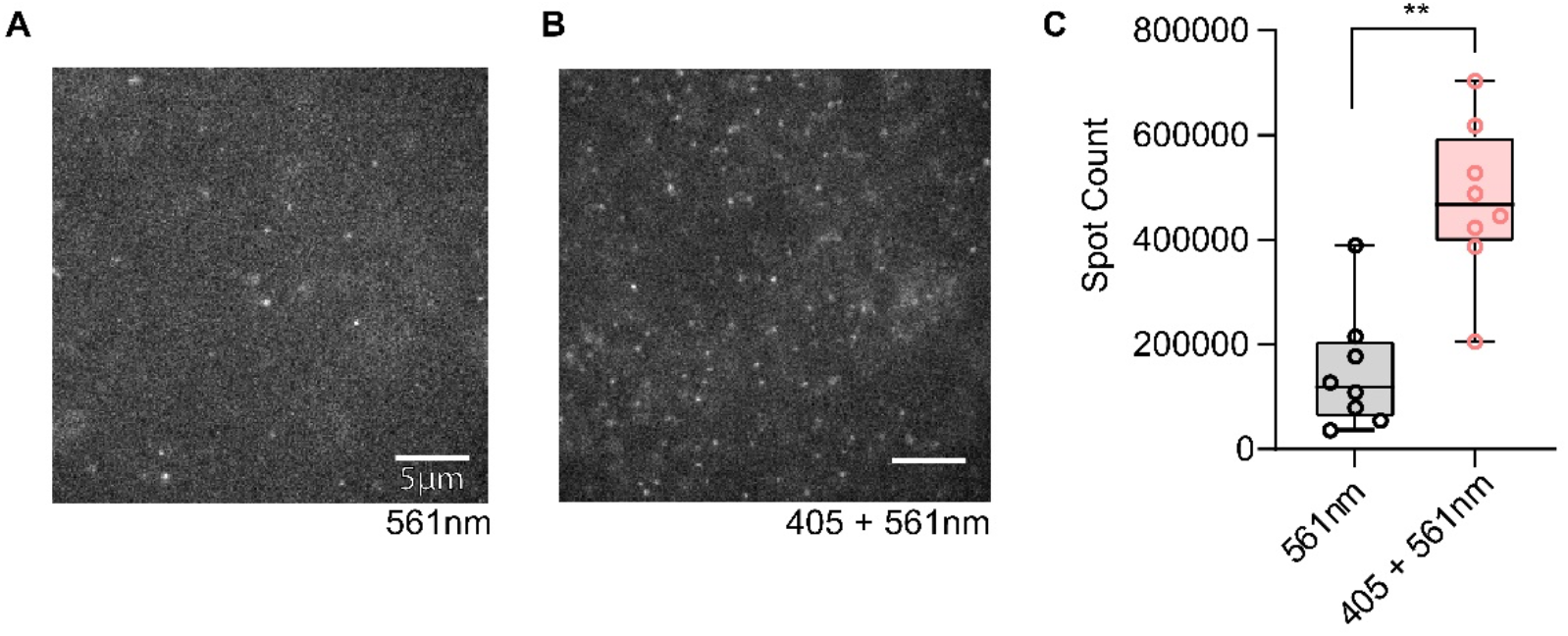
Photoconversion increases spot detection in the fly brain. (A) When recording in the fly brain for Sx1a-mEos2 molecules with only the 561 nm laser and (B) with a simultaneous UV-405 nm and 561 nm laser the total spot count (scale bars 5 µm). (C) shows a significant increase in spot detections for the 405-561 nm combination (n=8, *p*=0.0078, Spot Count CI 561 nm 53,883 - 245,102, Spot Count CI 405 + 561 nm 349,439 – 602,101, Wilcoxon test, data is ± 5-95^th^ percentile, scale bar is 5 µm). The relatively high number of spots in the 561 nm condition is most likely due to the high amount of auto-photoconversion after brain dissection under bright-field lights.

**Figure 1 – figure supplement 4:**
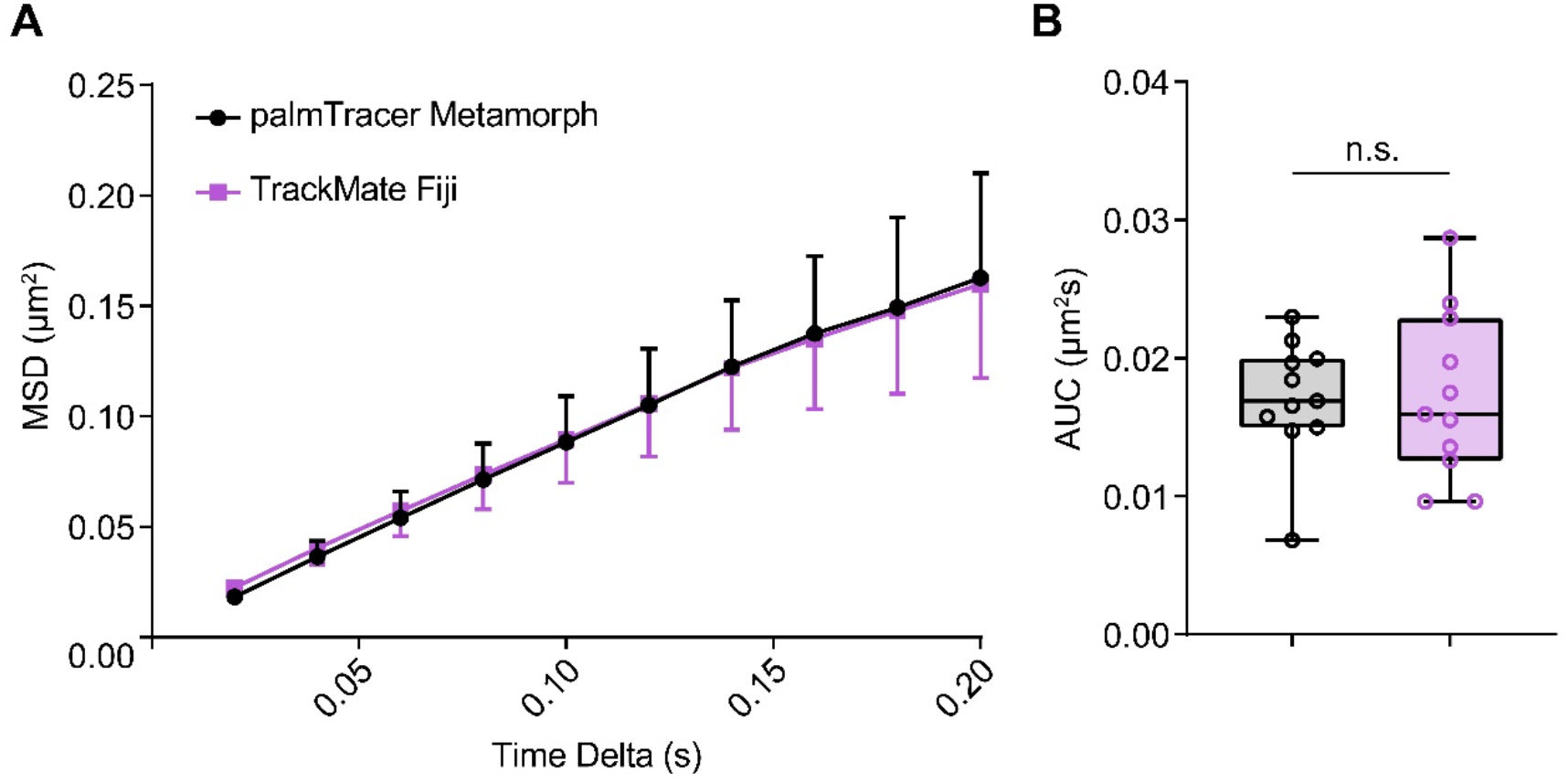
Validation of the semi-automated Single Particle Analysis (SPA) script employing TrackMate. To validate the SPA software that was used for all data analysis, we employed a known dataset that was analysed utilising the Metamorph plugin palmTracer. The data analysed were derived from rat pheochromocytoma PC12 cells that were transfected with a Munc18-1mEos2 (Kasula et al., 2016) tagged molecule. sptPALM was performed in the same way, except a lower exposure time of 20 msec was utilised. PC12 cells and Munc18-1mEos2 were provided by Fred Meunier, Queensland Brain Institute. (A) The MSD of Munc18-1mEos2 trajectories and (B) area under the curve (AUC) analysis reveals no significant difference between palmTracer and our custom TrackMate analysis scripts (n = 10, *p* = 0.898, Wilcoxon matched pairs signed rank test, 95% CI palmTracer 0.0142 – 0.0200, 95% CI TrackMate 0.0132 – 0.0213, MSD values presented as ± SD, AUC presented as ± 5-95 percentile).

**Figure 2 – figure supplement 1:**
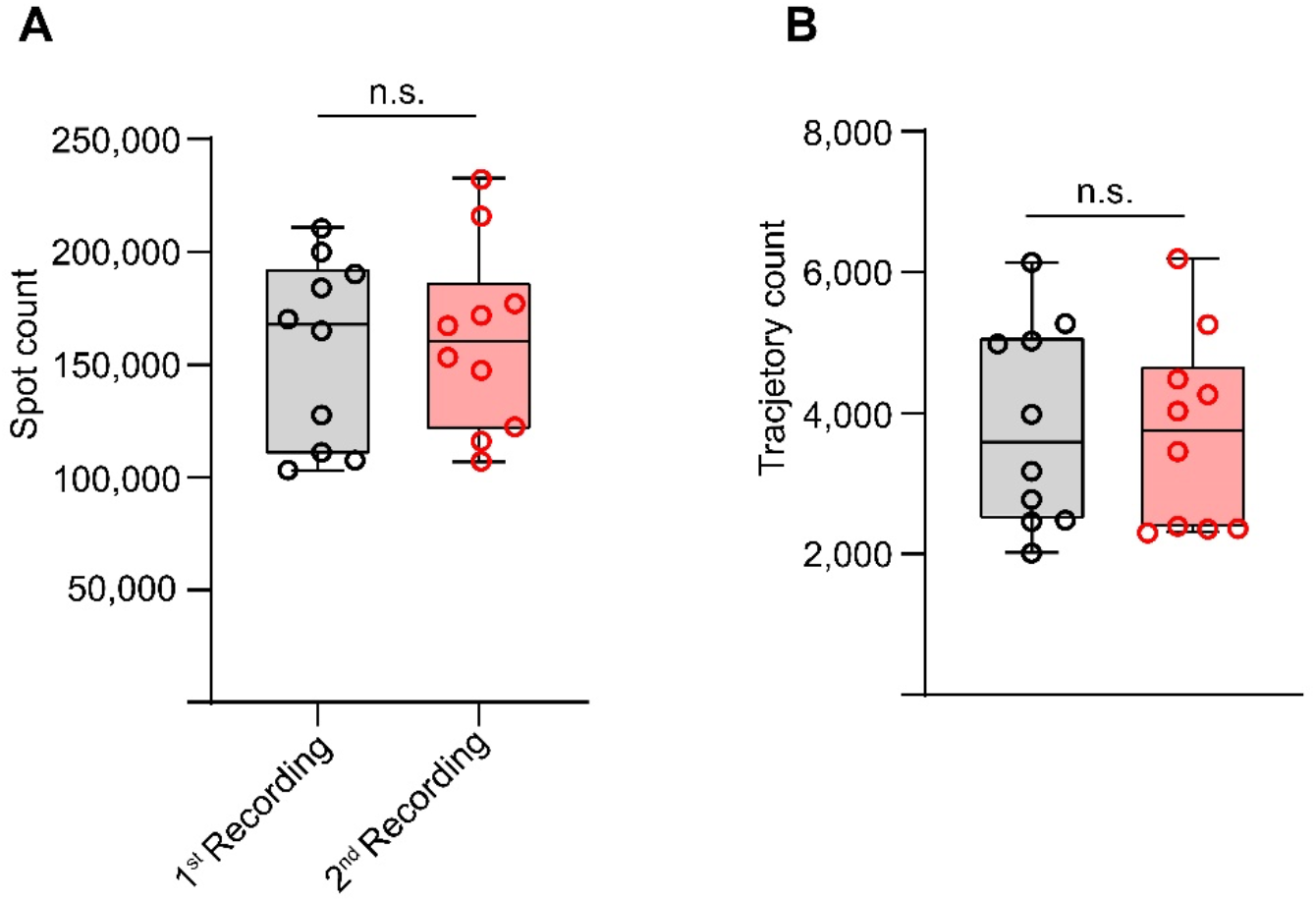
(A-B) Spot and trajectory counts between the first and second recording of Sx1a-mEos2 tracking experiments shows no difference in spot and trajectory counts (data from HL3.1 control recordings, n=10, *spot count p*=0.9118, *trajectory count p*=0.7394, n.s., not significant. Statistics performed for both with a Wilcoxon test, data is ± 5-95^th^ percentile).

**Figure 2 – figure supplement 2:**
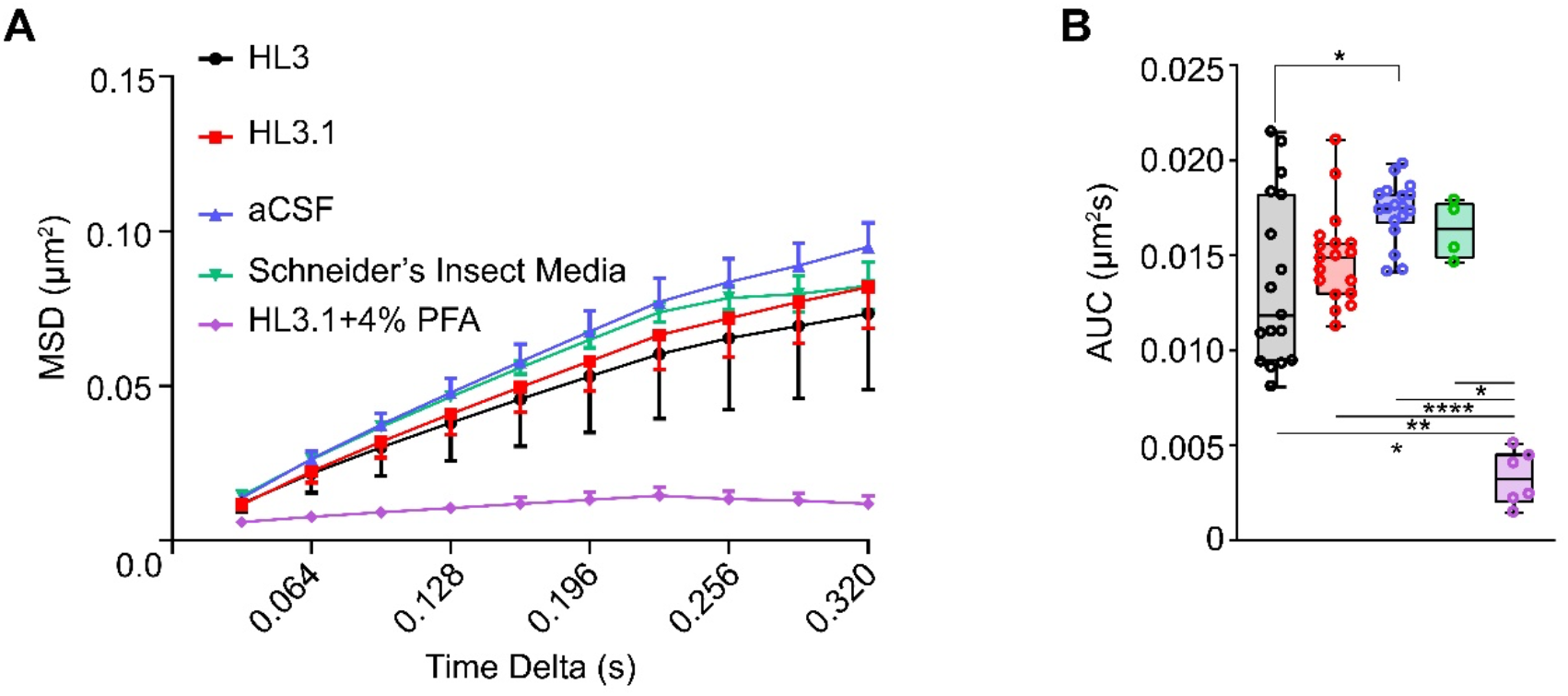
Comparison of imaging buffers on the mobility of Sx1a-mEos2 particles in the *Drosophila* brain. During method development phase, several imaging buffers were trialled for physiological relevance and consistency between samples. Three random brain regions were sampled in UAS-dTRPA1>R57c10-Gal4 flies at 30°C for stimulation in either hemolymph-like 3 (HL3), hemolymph-like 3.1 (HL3.1), artificial cerebrospinal fluid (aCSF), or Schneider’s insect media and compared for their consistency. Also included is a mobility control where brains were fixed in a 4% paraformaldehyde (PFA) before imaging in HL3.1 solution, to confirm that tracked molecules are not an artefact of the imaging buffer. (A) MSD curves for the average of the three imaging buffers utilised with (B) the AUC highlighting a significant difference between HL3 to aCSF (HL3 n=6, *p*=0.0058, AUC CI 0.0113 – 0.0160, HL3.1 n=6, *p*=0.0316, AUC CI 0.0137 – 0.0161, aCSF n=7, AUC CI 0.0165 – 0.0181, Schneider’s n=4, *p>*0.999, AUC CI 0.01383 – 0.01882, Kruskal-Wallis test, MSD data presented as ± SD, AUC data presented as ± 5-95 percentile). Despite aCSF providing the best consistency, HL3.1 was selected for its physiological relevance to *Drosophila* whilst retaining a degree of consistency above HL3. All imaging buffers were significantly different to the 4% PFA fixed brains, which showed minimal mobility effects (n=6, *p*=0.0212 HL3, *p*=0.0067 HL3.1, *p*<0.0001 aCSF, *p*=0.0116 Schneider’s, AUC CI 0.00174 – 0.00478, Kruskal-Wallis test).

**Figure 3 – figure supplement 1:**
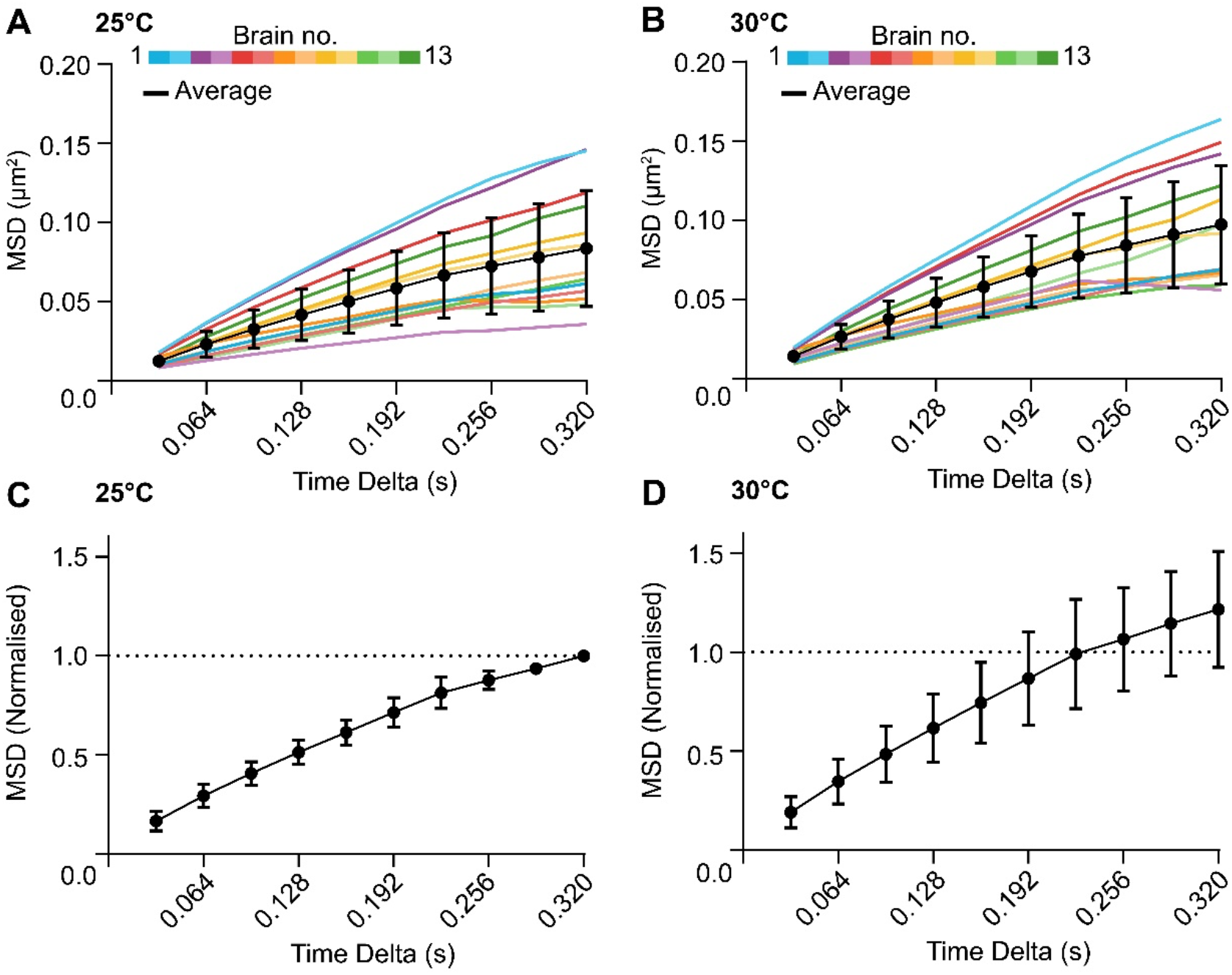
Normalization of neuronal stimulation MSD curves to baseline. (A) Raw and average MSD curves for Sx1a-mEos2 recorded in the adult fly brain at 25°C, each colour represents a different brain. (B) Raw and average MSD curves in the same brains as (A), but at 30°C for an internally controlled paradigm. (C) Normalised MSD curve for the raw data in (A). The peak value of the curve at time point 0.320 (s) was used to normalize each time point, such that the peak of the MSD curve at time 0.320 s is 1.0. (D) Normalized MSD curve for the raw data in (B). Each time point was normalized to the matched peak value of the corresponding baseline curves in (A). All average data is presented as ± SD.

**Figure 4 – figure supplement 1:**
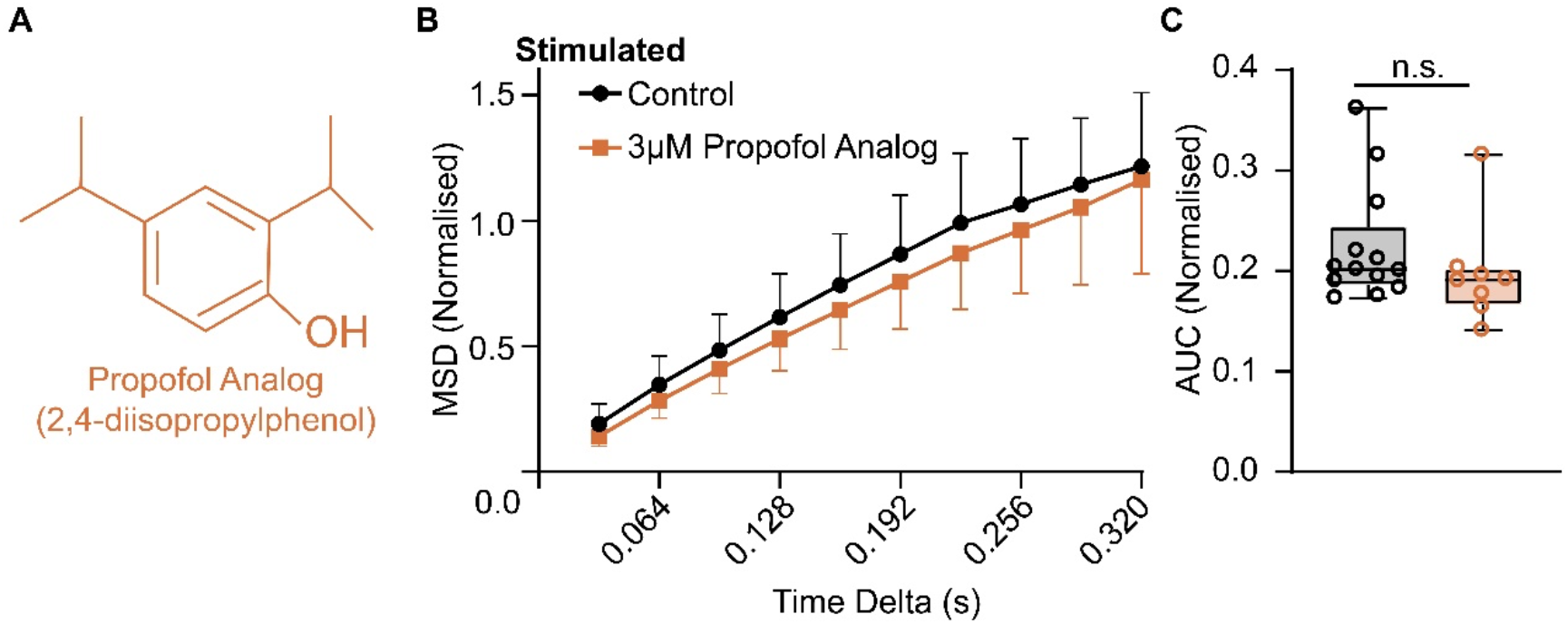
A structural propofol analog is not able to restrict Sx1a-mEos2 mobility. (A) Structure of the non-anaesthetic analog of propofol, (2,4-diisopropylphenol). Note the change in position of the hydroxyl group on the benzene ring from carbon 1 to carbon 3. (B) Under stimulation conditions, the MSD of Sx1a-mEos2 not able to be restricted in the presence of 3 µM of the propofol analog (orange) when compared to DMSO control (black), with no significant change in the (C) AUC (n=8, *p*=0.9866, AUC CI 0.1540 – 0.2406, Kruskal-Wallis test, data for MSD is ± S.D., data for AUC is ± 5-95^th^ percentile).

---

**Video S1:** Tracking individual Sx1a-mEos2 molecules in the fly brain.

**Video S2:** Absence of Sx1a-mEos2 mobility in fixed tissue.

**Video S3:** Absence of Sx1A-mEos2 tracking in HL3.1 solution.

**Video S4:** Conditional paralysis at 30°C in w^1118^;Sx1a-mEos2/+;UAS-TrpA1/R57C10-Gal4 flies and lack of paralysis at 30°C in w^1118^;Sx1a-mEos2/+;UAS-TrpA1/+ controls.

**Supplementary File 1:** Single Particle Analysis guide and associated software.

